# Carbon dots for efficient siRNA delivery and gene silencing in plants

**DOI:** 10.1101/722595

**Authors:** Steven. H. Schwartz, Bill Hendrix, Paul Hoffer, Rick A. Sanders, Wei Zheng

## Abstract

The Initiation of RNA interference (RNAi) by topically applied double stranded RNA (dsRNA) has potential applications for plant functional genomics, crop improvement and crop protection. The primary obstacle for the development of this technology is efficient delivery of RNAi effectors. The plant cell wall is a particularly challenging barrier to the delivery of macromolecules. Many of the transfection agents that are commonly used with animal cells produce nanocomplexes that are significantly larger than the size exclusion limit of the plant cell wall. Utilizing a class of very small nanoparticles called carbon dots, a method of delivering siRNA into the model plant *Nicotiana benthamiana* and tomato is described. Low-pressure spray application of these formulations with a spreading surfactant resulted in strong silencing of *GFP* transgenes in both species. The delivery efficacy of carbon dot formulations was also demonstrated by silencing endogenous genes that encode two sub-units of magnesium chelatase, an enzyme necessary for chlorophyll synthesis. The strong visible phenotypes observed with the carbon dot facilitated delivery were validated by measuring significant reductions in the target gene transcript and/or protein levels. Methods for the delivery of RNAi effectors into plants, such as the carbon dot formulations described here, could become valuable tools for gene silencing in plants with practical applications in plant functional genomics and agriculture.

## INTRODUCTION

RNAi is composed of inter-related pathways that mediate transcriptional gene silencing (TGS) via methylation of genomic sequences, post-transcriptional gene silencing via the cleavage of targeted RNA sequences, or translational repression of targeted transcripts (Matzke and Matzke, 2004, Frizzi and Huang, 2010). In plants, these pathways have a role in resistance to pathogens (Rosa *et al*., 2018a) and are also required for normal development (Carrington and Ambros, 2003). The use of RNAi to silence specific genes has been a valuable tool in plant functional genomics (McGinnis, 2010, Kumar and Salar, 2017). There have also been a number of applied biotechnology applications of RNAi (Frizzi and Huang, 2010). This includes improving the nutritional composition of crops (Huang *et al*., 2004, Mroczka *et al*., 2010, Chawla *et al*., 2012) and providing resistance to various plant pathogens that can significantly reduce crop yield and quality (Rosa *et al*., 2018b). Applications for the control of insect pests have also been developed (Baum *et al*., 2007).

The first uses of gene silencing in plant biology involved stable transformation with “antisense" or “co-suppression” constructs. With a better understanding of the RNAi pathway, more efficient methods were developed that utilize dsRNA hairpins (Smith *et al*., 2000) and artificial microRNAs (Schwab *et al*., 2006). Another important advancement in RNAi technology has been the development of methods for transient gene silencing, such as virus-induced gene silencing (VIGS) (Burch-Smith *et al*., 2004, Watson *et al*., 2005, Becker and Lange, 2010) and *Agrobacterium* infiltration (Johansen and Carrington, 2001). These transient silencing systems allow for the rapid testing of gene function. Both VIGS and Agrobacterium infiltration require the preparation of constructs for the expression of the RNAi effector and containment of the pathogen infected plants. Efficient delivery of topically applied RNAi effectors would be another valuable tool for plant functional genomics and may have some practical applications in agriculture.

Because of the potential for therapeutic applications, delivery of RNAi effectors has been extensively studied in animal systems. Many nanoparticle-based transfection agents that enhance delivery into animal cells have been described (Kozielski *et al*., 2013). Common classes of these transfection agents include lipid nanoparticles, cationic polymers, cell-penetrating peptides, and inorganic nanoparticles. The nanocomplexes that are formed by the interaction of these transfection agents with nucleic acids provide some protection from nucleases and facilitate cellular uptake by endocytosis or membrane fusion. There have been a number of reports describing the use of transfection agents for delivery to plant cells (Unnamalai *et al*., 2004, Cheon *et al*., 2009, Eggenberger *et al*., 2011, Lakshmanan *et al*., 2013, Numata *et al*., 2014, Ziemienowicz *et al*., 2015, Kimura *et al*., 2017, Golestanipour *et al*., 2018, Miyamoto *et al*., 2019). The efficiency of many transfection agents, however, may be limited by delivery barriers that are unique to plants (Figure 1). The plant cell wall is a particularly challenging barrier for the delivery of RNA or other macromolecules. The dense polysaccharide matrix of the cell wall has a size exclusion limit that is between 3 and 10 nm in diameter (Carpita *et al*., 1979, Baron-Epel *et al*., 1988, Carpita and Gibeaut, 1993). While many of the nanocomplexes used to transfect nucleic acids, have a size in the range of 100 to 200 nm. This is 10 to 20-fold larger than the size exclusion of the plant cell wall. Recently, there have been several reports of using smaller nanostructures such as single-walled carbon nanotubes (Demirer *et al*., 2019, Kwak *et al*., 2019) and DNA nanoparticles (Zhang *et al*., 2019) for delivery of nucleic acids into plant cells.

**Figure 1.**
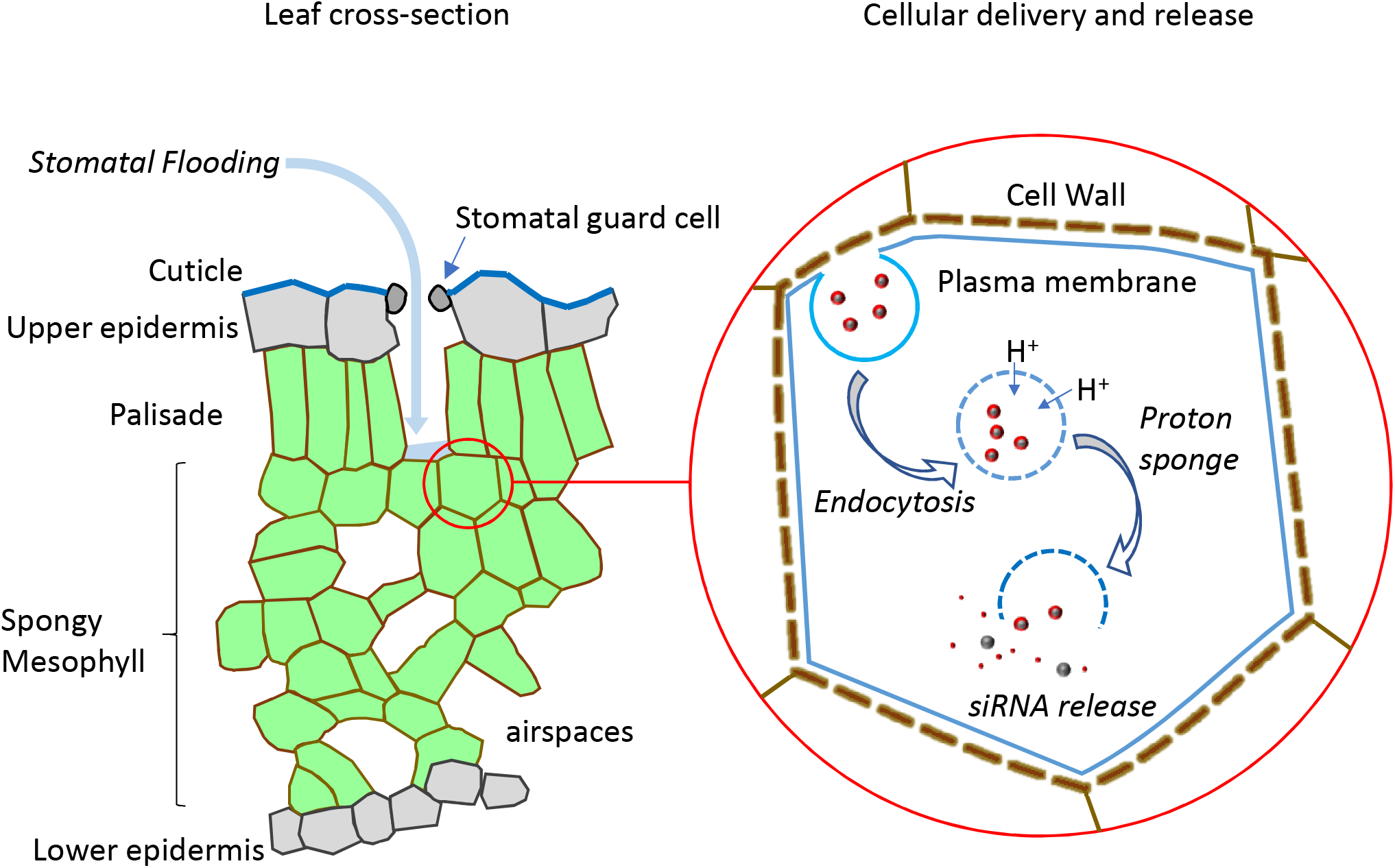
Strategy for topical RNAi in plants. The first barrier for delivery is the cuticle, a water impermeable layer, that covers all above ground parts of the plant. Stomates, the pores that allow for gas exchange, can be a point of entry for formulations. With a spreading surfactant, formulations can flow across the leaf surface and “flood” stomates for delivery to the mesophyll cells. Once in the leaf apoplast, nanocomplexes containing the siRNA cargo would need to diffuse through the cell wall to reach the plasma membrane. The relatively small size exclusion limit of the cell wall (< 10 nm) would likely restrict the movement of larger nanocomplexes. Smaller nanocomplexes are able to reach the plasma membrane and facilitate cellular uptake by endocytosis. Escape from the endomembrane vesicles is achieved by a phenomenon called the “proton-sponge” effect, which causes osmotic swelling and lysis of the vesicles (Behr, 1997). The siRNA cargo is now accessible to the RNAi machinery. For post-transcriptional gene silencing (PTGS), the guide strand is loaded into the RISC complex allowing for specific cleavage of mRNAs with complementary sequences.

There are additional classes of nanoparticles that may be well suited for plant delivery. Quantum dots are very small nanoparticles that have shown good efficacy in the transfection of animal cells (Yezhelyev *et al*., 2008). A significant disadvantage to quantum dots is that they are most often made from heavy metals. In recent years, carbon dots have received a considerable amount of attention as a “green” alternative to the heavy metal containing quantum dots (Reckmeier *et al*., 2016, Yao *et al*., 2019). Much of the interest in carbon dots has evolved around their optical properties, which have practical applications for bioimaging, photocatalysis, photovoltaic cells, and in light-emitting diodes. There are reports on the use of carbon dots for transfection of plasmids, long dsRNA, and siRNA into animal cells (Liu *et al*., 2012, Wang *et al*., 2014, Das *et al*., 2015, Pierrat *et al*., 2015). The uptake of carbon dots in plants has been studied (Li *et al*., 2016, Li *et al*., 2018, Qian *et al*., 2018), but their use for delivery of nucleic acids into plant cells has not yet been reported. Because of their small size, it was hypothesized that carbon dots may be useful for the delivering dsRNA through the plant cell wall and subsequent barriers.

## RESULTS

### Preparation and purification of carbon dots

Carbon dots can be synthesized by a “top down” approach, which involves decomposition of structured carbon precursors such as graphene. Alternatively, the “bottom-up” approach begins with carbonization of simple carbon precursors such as organic acids, sugars, or amino acids (Yao *et al*., 2019). The bottom-up methods for producing carbon dots usually involve a hydrothermal reaction or pyrolysis of the carbon precursor to produce nanoparticles with sizes that are typically between 1 and 10 nm. Surface functionalization/passivation of the carbon dots can increase the colloidal stability and allow for the binding of various ligands. Functionalization with amines produces particles with a positive charge, that can bind with the negative charges of the phosphate backbone in nucleic acids. Polyethyleneimine (PEI) is a commonly used cationic polymer for the functionalization of carbon dots. Citrate derived carbon dots that have been functionalized with branched PEIs (bPEIs) have been used to deliver plasmid DNA and siRNA to animal cells (Liu *et al*., 2012, Pierrat *et al*., 2015).

Functionalization of carbon dots with PEI is often done in a “one-pot” synthesis where carbonization of the precursor and functionalization occur at the same time. This is possible because aqueous solutions of PEI are relatively thermostable. It has been reported that H_2_O_2_ can assist in the formation of carbon dots directly from PEI (Zhou *et al*., 2015, Wang *et al*., 2017). In these examples, PEI served as both the carbon source for the core and the nitrogen source for surface passivation. In the work described here, carbon dots were produced directly from bPEIs of various molecular weights by heating solutions in a mixture of chloroform and methanol to 155 °C for a relatively short time. The product of these reactions displayed the characteristic color development of carbon dots. The PEI derived carbon dots had an absorption max of 363 nm and produced a characteristic blue fluorescence with an emission maximum at 460 nm (Figure S1).

Carbon dot preparations may contain a heterogenous mixture of precursors, by-products, and nanoparticles with different sizes and physical properties. Therefore, the preparations were fractionated by size exclusion chromatography (Figure 2A) and the size distribution of the nanoparticles within the primary peaks was determined by dynamic light scattering (Figure 2B). Based on the size exclusion chromatography and dynamic light scattering measurements, the relative size of the carbon dots correlates well with molecular weight of the PEI precursors. The smallest PEIs used in this study, with average molecular weights of 1200 and 1800, produced the smallest carbon dots. The carbon dots produced from 5 KD, 10 KD, or 25 KD bPEIs were progressively larger.

**Figure 2.**
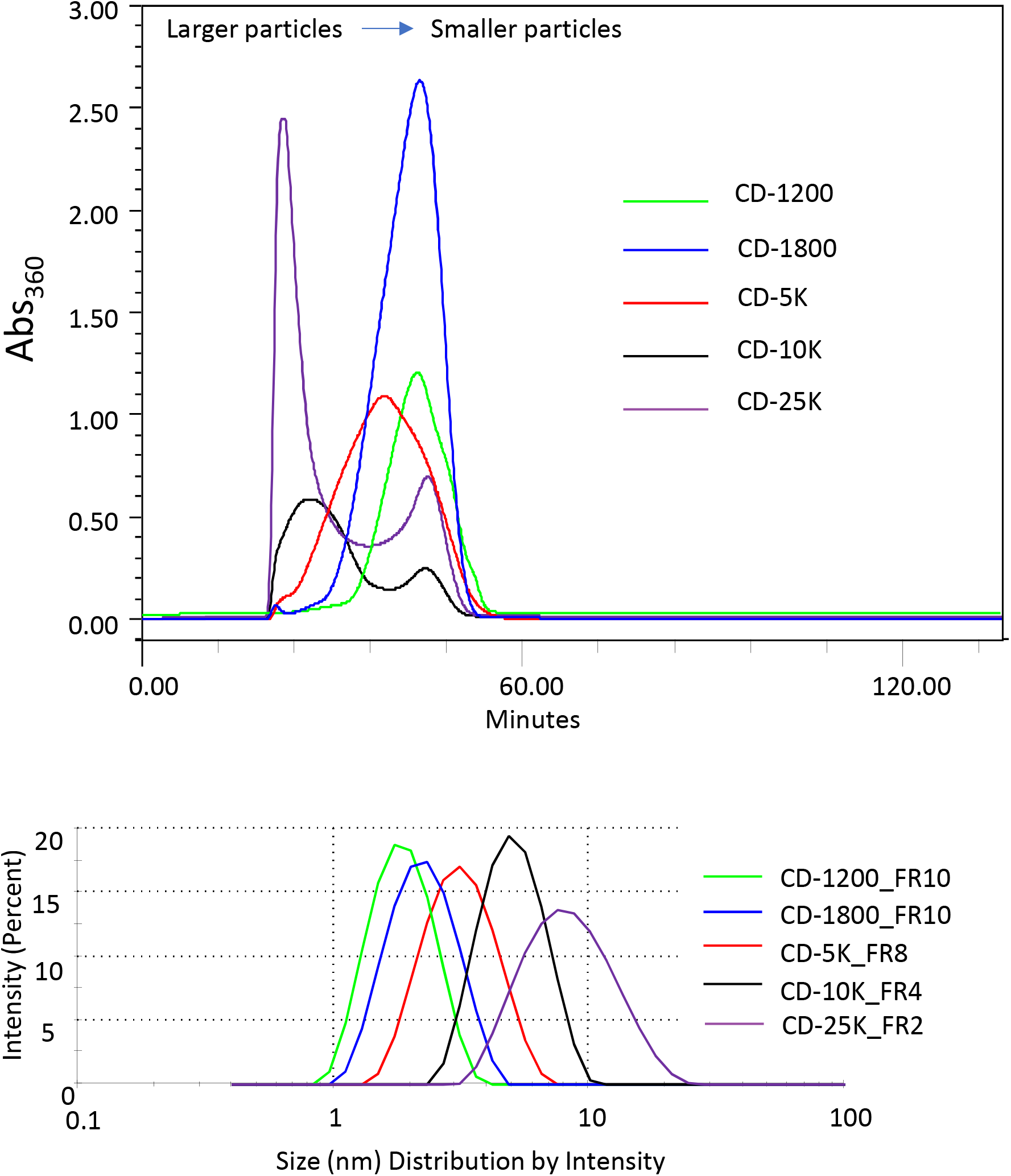
Purification and size characterization of carbon dots. *(A)* Carbon dots were produced from different molecular weight polyethyleneimines and purified by size exclusion chromatography. The elution of the carbon dot preparations was monitored at 360 nm. *(B)* The particle size distribution within the purified fractions was determined by dynamic light scattering (DLS).

### Carbon dots provide protection form nucleases

The binding of a 124-base pair dsRNA to carbon dots and protection from nuclease activity was demonstrated by agarose gel electrophoresis (Figure 3). The absence of a band after formulation with carbon dots indicates binding of dsRNA, which can subsequently be released by the addition of sodium dodecyl sulfate (SDS). Resistance to nucleases was tested by treatment of dsRNA alone or dsRNA bound to carbon dots with RNase III from *E. coli*. In assays with the dsRNA alone, significant degradation was observed after one minute and the dsRNA was almost completely degraded by 15 minutes. While the dsRNA that was formulated with the carbon dots, was still intact after a 60-minute incubation with RNase III. The enhanced resistance to nucleases could significantly enhance efficacy in plants, which contain high levels of RNase activity in the extracellular apoplast (Sangaev *et al*., 2011).

**Figure 3.**
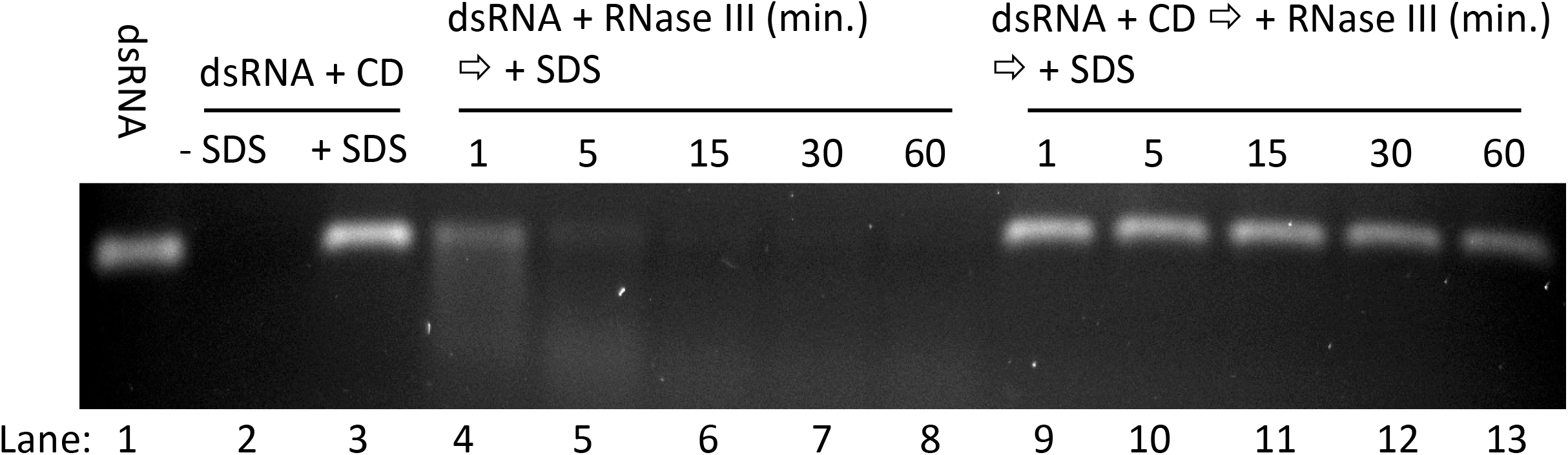
Carbon dots can bind dsRNA and provide protection from nucleases. Agarose gel electrophoresis with dsRNA alone or dsRNA formulated with carbon dots (lanes 1 and 2). Formulation with the carbon dots prevents migration into the gel and binding of ethidium bromide. After binding to carbon dots, dsRNA can be released by the addition of SDS (lane 3). Rnase III was incubated with naked dsRNA (lanes 4-8) or dsRNA that was bound to carbon dots (lanes 9-13) for the indicated times to determine if carbon dots can provide protection from degradation. Prior to electrophoresis, SDS was added to release the dsRNA from the carbon dots.

### Carbon dot formulations are efficacious in silencing *GFP* in the 16C line of *N. benthamiana*

The Green Fluorescent Protein (GFP) expressing 16C line of *Nicotiana benthamiana* is an often-used model for the study of RNAi (Ruiz *et al*., 1998, Dalakouras *et al*., 2016, Bally *et al*., 2018). In wild type plants, chlorophyll in the leaves displays a strong red fluorescence under UV or blue lights. In the GFP transgenic lines, this red fluorescence is masked by the green fluorescence. Silencing of the *GFP* is then easily detected by the un-masking of the red chlorophyll fluorescence. For the silencing experiments in this study, 22-mer siRNAs were used. Several lines of research have demonstrated that the initiation of silencing with 22-mer siRNAs may result in greater secondary siRNA production, which can enhance RNAi phenotypes (Mlotshwa *et al*., 2008, Dalakouras *et al*., 2016, Taochy *et al*., 2017, Hendrix *et al*., Manuscript in preparation). The sequence of the *GFP* targeting 22-mer and other siRNAs used in this study are shown in Table S1.

True leaves 3 and 4 from 17-day old plants (Figure S2) were treated by adaxial spray of formulations with 0.4% BREAK-THRU^®^ S 279, a non-ionic spreading surfactant. This allows entry of the formulation into plants by stomatal flooding, which is a commonly used method for delivery of agrochemicals into leaf tissue. A typical concentration of siRNA used was 12 ng/μL with an estimated application volume of 3.8 μL/cm^2^. These application parameters give an approximate siRNA use rate of 45 ng/cm^2^. In the initial screen for efficacy, carbon dots produced from different molecular weights PEIs were tested (Figure 4). With the lowest molecular weight PEI precursors (Mw 1200 and 1800), there was little or no visible GFP silencing. A limited amount of the silencing was observed for carbon dots derived from the largest PEI tested (Mw 25 KD). Much higher levels of silencing were observed with the carbon dots that were produced from the 5 KD bPEI or the 10 KD bPEI.

**Figure 4.**
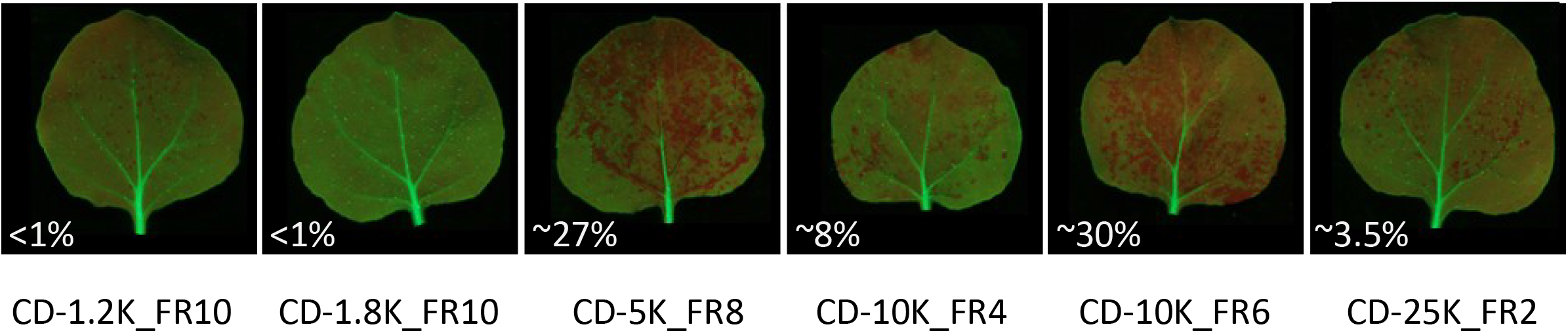
GFP silencing observed with carbon dots produced from different molecular weight PEIs. Silencing efficacy of fractions corresponding to the primary peaks from each of the different molecular PEI precursors were tested for activity with a 22-mer targeting *GFP* at a concentration of 12 ng/μL. At five days after treatment, the GFP Fluorescence was visualized under a strong blue light. The appearance of red chlorophyll fluorescence under the blue lights is indicative of GFP silencing. Representative leaf images for each of the carbon dot preparations are shown in the figure. The percent silencing area below each picture was measured digitally with 4 leaves for each treatment. Fraction #6 from the 10 KD bPEI preparation, which eluted after the primary peak was included in the analysis, because it displayed better silencing activity than the fraction from the primary peak.

Because unmodified PEIs are one of the most often-used transfection agents for delivery to animal cells, the unmodified 5 KD bPEI was also tested for delivery and silencing efficacy with the 16C line (Figure S3). With these buffer conditions, the unmodified 5 KD bPEI was not an effective transfection agent for delivery to intact plant cells. Improved delivery efficiency can be achieved with unmodified PEIs when a high concentration of an osmoticum is included in the formulations(Zheng *et al*., Manuscript in preparation).

In the initial screen, some differences were observed in the delivery efficacy of fractions from the same preparation. Fraction 6 from the 10 KD preparation, for example, displayed much higher silencing efficacy than fraction 4 from the same preparation (Figure 4). As part of the optimization process, additional fractions from the 5 KD bPEI preparation (CD-5K) were tested for efficacy in silencing *GFP* in the 16C line. Based on the image analysis, fraction #5 from the CD-5K preparation was the most efficacious (Table 1 and Figure S4). Later fractions, which contain smaller carbon dots, were less efficacious in silencing *GFP*. While a small size may be a prerequisite for delivery through the plant cell wall, it does not appear to be the only factor affecting delivery and silencing efficiency. The particle characteristics optimal for movement through the cell wall may not be optimal for other barriers to delivery. Endocytosis of nanoparticles in animal cells can be affected by size, surface chemistry, shape, and rigidity (Zhang *et al*., 2015).

**Table 1.** Percent silenced area with different fractions of CD-5K. Different fractions of the CD-5K preparation were formulated with the *GFP* targeting 22-mer at a concentration of 8 ng/μL. The percent silencing relative area relative to the entire leaf area was calculated digitally. The connected letter report displays the percent silencing and a statistical comparison of silenced area produced with the different fractions. In the LS-Means Student’s t, the fractions not connected by a letter are statistically significant at a p-value of 0.05. The images used for the analysis are included as Figure S4.

| Fraction |  |  |  | Silenced area<br>(Least Sq Mean) |
| --- | --- | --- | --- | --- |
| Fraction #5 | A |  |  | 43.56 |
| Fraction #6 | A |  |  | 36.19 |
| Fraction #7 | A | B |  | 35.60 |
| Fraction #8 | A | B |  | 28.48 |
| Fraction #9 |  | B |  | 20.35 |
| Fraction #10 |  |  | C | 4.89 |

For molecular validation of gene silencing, the CD-5K_FR5 was used (Figure 5). Application of a carbon dot formulation with a non-target siRNA did not impact the typical green fluorescence of the 16C line under blue lights. While plants that were sprayed with a formulation containing a *GFP* targeting 22-mer siRNA displayed a strong red fluorescence that covered most of the leaf. The level of *GFP* transcript reduction, measured by qRT-PCR, was 84%. Western analysis showed a similar reduction in GFP protein levels. This level of *GFP* silencing was sufficient to initiate systemic spread of silencing. By 12 days after treatment, *GFP* silencing in newly emerging leaves became apparent (Figure S5).

**Figure 5.**
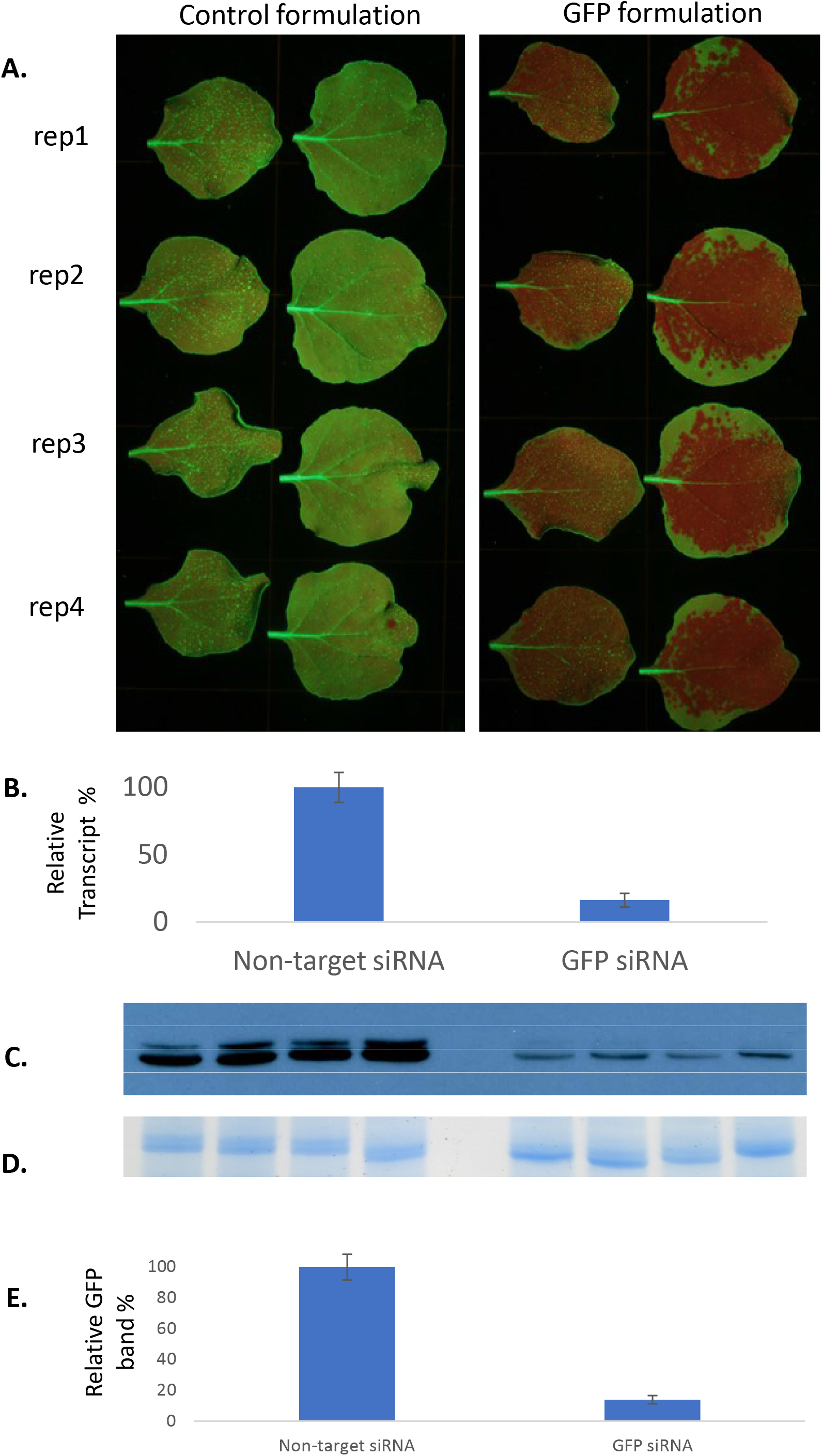
Molecular validation of *GFP* silencing in the 16C line on *N. benthamiana*. Leaves 3 and 4 were sprayed with carbon dot formulations containing a non-target siRNA (NT) or a 22-mer siRNA targeting the *GFP* transgene. The final concentration of siRNAs in the formulations was 12 ng/μL. *(A)* Silencing of *GFP* was monitored by fluorescence under blue lights at 5 days after treatment. *(B)* The silencing of *GFP* was validated by qRT-PCR analysis of transcript levels P-value=0.00048. *(C)* Western blot of GFP protein. *(D)* For the Coomassie stained image, the same protein extracts were run on a separate gel to demonstrate similar loading. *(E)* Quantification of band intensity with ImageJ. Error bars are standard error of the mean (SEM). The reduction in GFP protein was statistically significant with a P-Value=0.0000651

### Carbon dot formulations are efficacious in silencing endogenous genes in *N. benthamiana*

The silencing of endogenous genes was tested by targeting genes encoding the H and I sub-units of Magnesium Chelatase (MgChe), an enzyme necessary for chlorophyll synthesis. True leaves 3 and 4 from 17-day old *N. benthamiana* plants were sprayed with carbon dot formulations containing a nontarget siRNA, an siRNA targeting *MgCheH*, or an siRNA targeting *MgCheI* (Figure 6A). Leaves that were sprayed with formulations targeting either the *H* or *I* sub-units, displayed spots and patches of bleaching that is indicative of reduced chlorophyll accumulation. Over several experiments, the bleaching phenotype was strongest on the younger leaf four and with the formulation targeting the *MgCheH* gene. Analysis of *MgCheH* transcript levels by qRT-PCR showed a 79% reduction in the phenotypic tissues at five days after treatment (Figure 6B). The bleaching phenotype persisted for the duration of the experiments on the application leaves, which was up to 20 days after treatment (Figure S6).

**Figure 6.**
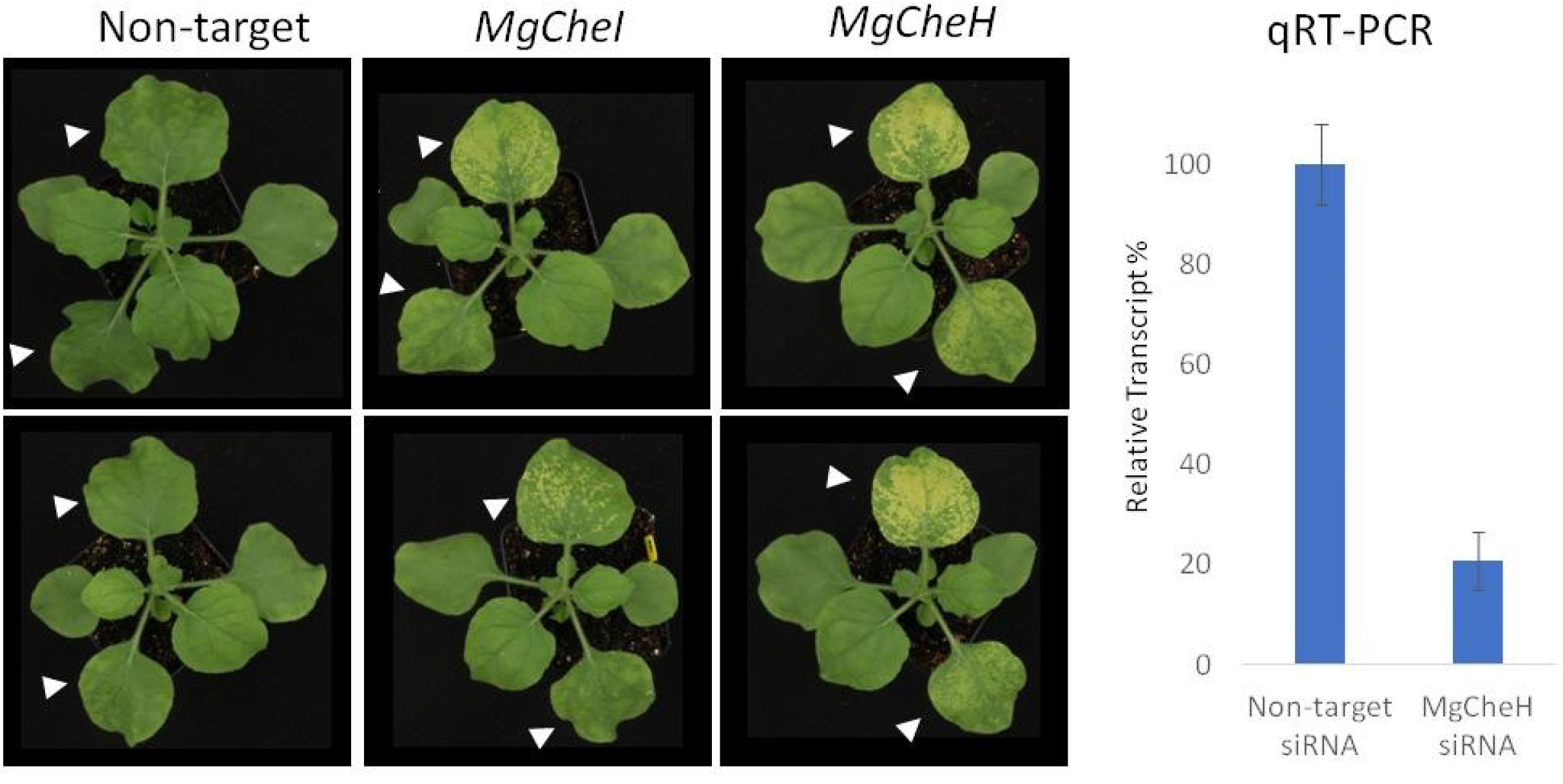
Silencing of the *Magnesium Chelatase* (*MgChe*) *H* or *I* sub-units in *N. benthamiana*. *(A)* Bleaching phenotype in treated leaves. Leaves 3 and 4 were sprayed with carbon dot formulations containing a non-target siRNA, a 22-mer targeting *MgCheI*, or a 22-mer targeting *MgCheH*. The final concentration of siRNAs in the formulations was 12 ng/μL. Images were taken 4 days after treatment. *(B)* Transcript analysis of *MgCheH*. At five days after treatment, the tissue was sampled for qRT-PCR analysis. Error bars are SEM. The reduction in *MgCheH* transcript was statistically significant with a with a P-Value=3.48E-06

The *MgCheH* silencing phenotype was used to evaluate the stability of carbon dot formulations. A single batch of formulation with the *MgCheH* targeting 22-mer was sprayed on plants 1 day, 1 week, or 2 weeks after preparation. Plants were imaged 5 days after treatment (Figure S7). With the spray application at one week, there did not appear to be a significant loss in efficacy of the formulation. After two weeks of storage at room temperature, there appears to be only a small reduction in efficacy of the formulation; indicating that the siRNA is mostly intact, and that the formulation has good colloidal stability.

### Carbon dot formulations are efficacious for siRNA delivery and silencing in tomato

The efficacy of carbon dot formulations for siRNA delivery and silencing in tomato was tested with a line expressing an enhanced Green Fluorescent Protein (eGFP). Plants treated with a formulation containing a non-target siRNA displayed a strong green fluorescence (Figure 7A). With formulations containing a *eGFP* targeting 22-mer, silencing is apparent as spots and patches at the lower siRNA concentrations (2 and 4 ng/μL). At an siRNA concentration of 8 ng/μL, the silencing phenotype covers most of the leaf area. By Western blot analysis of whole leaf extracts, an 88% reduction in GFP protein levels was observed (Figure 7B and 7D).

**Figure 7.**
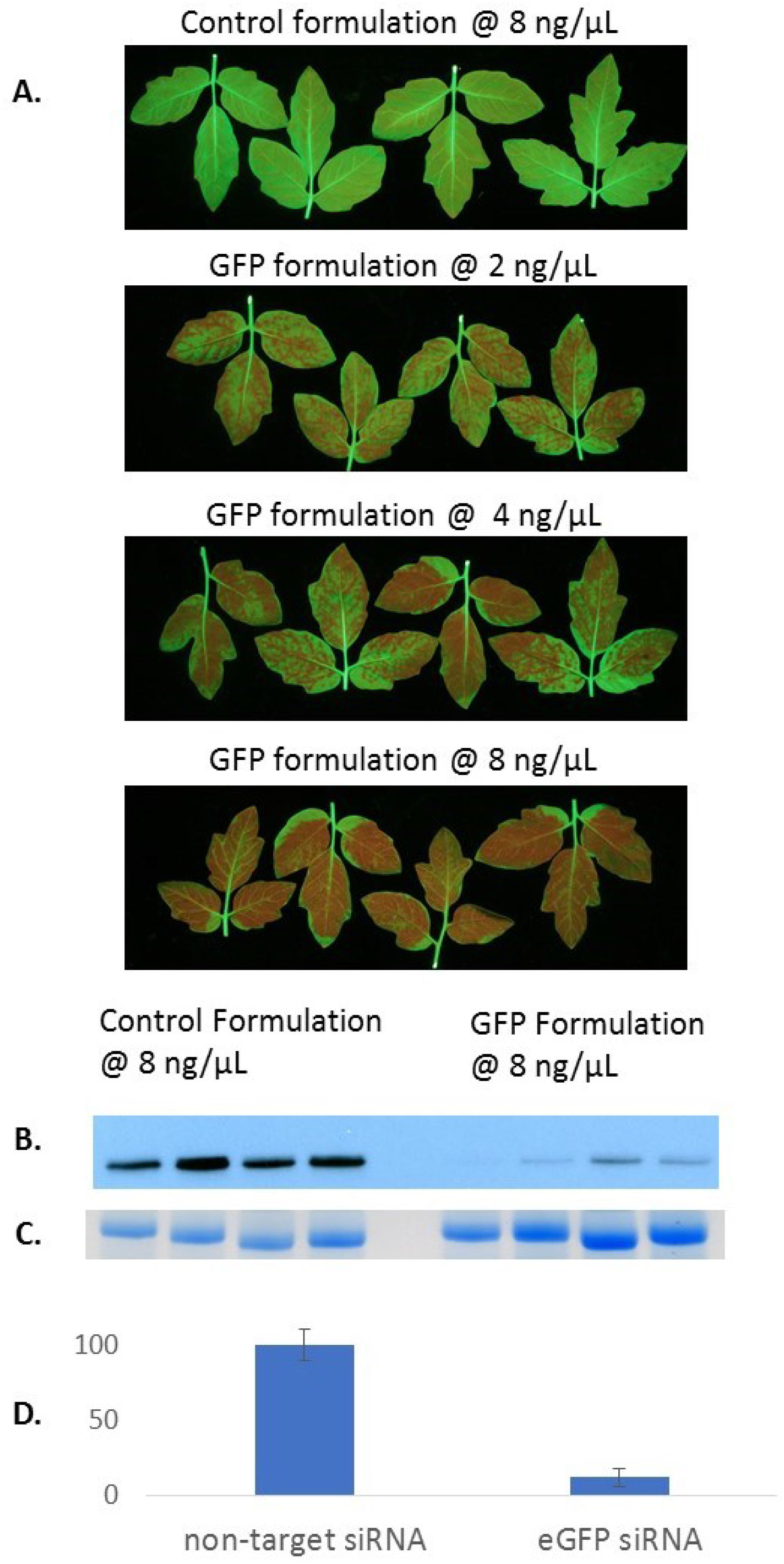
GFP silencing in fully expanded tomato leaves. True leaves 3 and 4 received an abaxial spray of carbon dot formulations containing a non-target siRNA or a 22-mer siRNA targeting *GFP*. The final concentration of siRNAs is shown on the figure. *(A)* The GFP fluorescence was imaged at 5 days after treatment. *(B)* Western blot analysis of GFP protein levels from whole leaf extracts. *(C)* Coomassie stained gel to demonstrate equal loading. *(D)* Quantification of band intensity with ImageJ. Error bars are SEM. The reduction in GFP protein was statistically significant with a P-Value =0.00041

Stomatal flooding works well with fully expanded leaves. In younger leaves, spray application of carbon dot formulations results in strong silencing towards the tip with more limited silencing at the base (Figure 8). In basipetal plants, such as tomato, functional stomates develop first at the leaf tip. The limited silencing at the base of the young leaves is likely due to reduced flooding where stomates have not yet fully developed. With the limitations in stomatal flooding, other methods maybe more amenable for silencing genes in very young tissues. Delivery methods for DNA or RNA that rely on physical disruption of barriers have also been used (Shang *et al*., 2007, Dalakouras *et al*., 2016). These physical delivery methods appear to be most effective with young tissues and could be complementary to the method described here.

**Figure 8.**
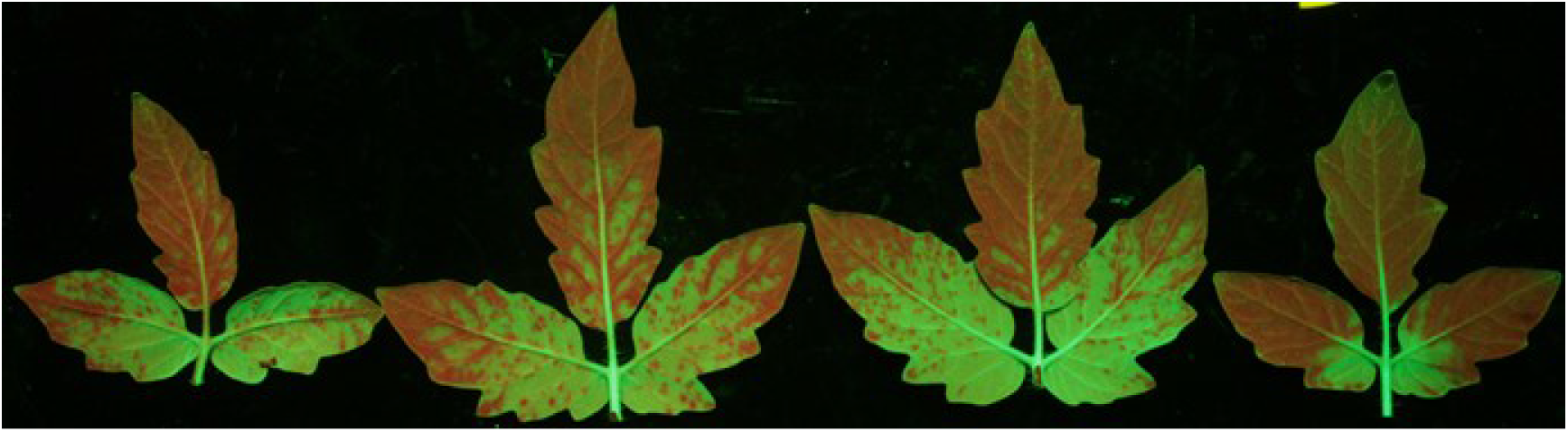
Silencing in younger tomato leaves. Fluorescence image of tomato leaves 5 days after treatment with a carbon dot formulation containing the *eGFP* targeting 22-mer at 8 ng/μL.

### An alternative method for production of carbon dots

Given the high level of activity observed with CD-5K, a method was developed to produce carbon dots from glucose with 5 KD bPEI functionalization in a “one-pot” synthesis. These carbon dots, which were produced at a lower temperature and in an aqueous solution, showed the same high levels of activity for the silencing of target genes in *N. benthamiana* and tomato (Figure 9). With the lower temperature and the aqueous solvent used for production of the glucose carbon dots, methods could be developed for their production that would require less specialized lab equipment.

**Figure 9.**
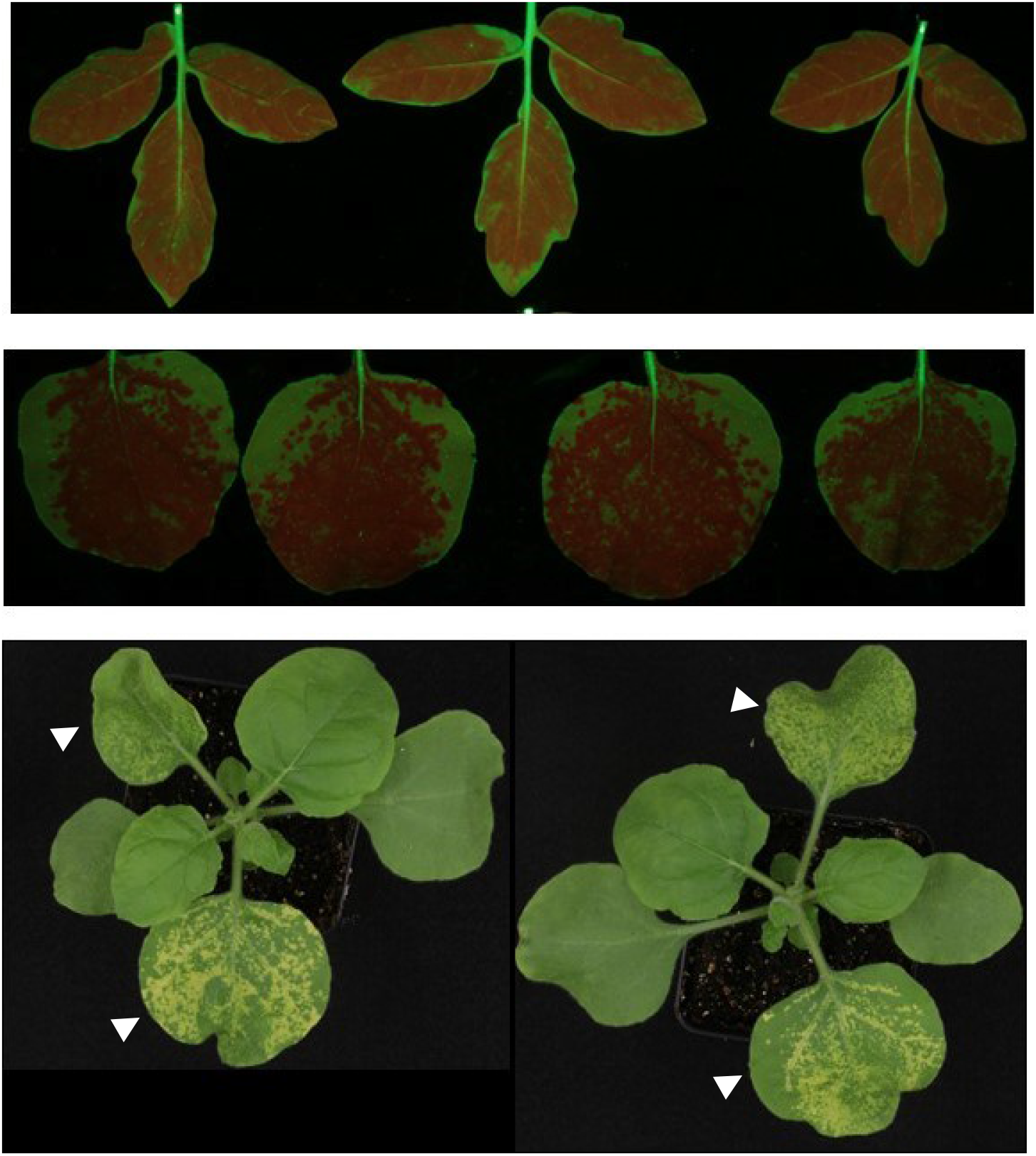
Efficacy of 5K bPEI functionalized glucose-carbon dots. *(A) GFP* silencing in tomato *(B) GFP* silencing in the 16C line of *N. benthamiana (C) MgCheH* silencing in *N. benthamiana*

## DISCUSSION

Transient gene silencing methods, such as VIGS or Agrobacterium infiltration, have been important tools for studying the functions of plant genes. The development of delivery methods for topically applied siRNA would be another valuable tool in the RNAi toolbox. In addition to simplicity, topical dsRNA methods would not require containment of the treated plants. As discussed in the introduction, transfection agents that are used in animal systems may have limited efficacy in plants due to the cell wall size limitations. Recently, there have been several reports describing the use of nanostructures for delivery of nucleic acids into plant cells (Mitter *et al*., 2017, Demirer *et al*., 2019, Kwak *et al*., 2019, Zhang *et al*., 2019). Another class of small nanoparticles called carbon dots have been used in animal systems for bio-imaging, delivery of various drugs, and the delivery of nucleic acids. Based upon their small size, it was hypothesized that carbon dots may readily pass through the cell wall and provide an effective delivery method for plant cells.

With a short reaction time and moderate temperatures, carbon dots could be produced directly from bPEIs in a solution of chloroform and methanol. An alternative method of producing carbons dots from glucose and bPEIs in an aqueous solution and at a lower temperature is also described. Amongst the PEI precursors used, carbon dots produced from the 5 KD bPEI displayed the highest level of activity for delivery. Purification of the carbon dots by size exclusion chromatography was an important step in the optimization. Components of a preparation that can bind RNA but are less efficacious for delivery would likely reduce silencing activity by sequestering the siRNA. With a low-pressure spray application, carbon dot formulations showed good efficacy in the delivery of siRNA to *N. benthamiana* and tomato leaves. Strong visible leaf phenotypes were observed when silencing *GFP* transgenes or endogenous genes necessary for chlorophyll synthesis. This gene specific silencing was validated by molecular characterization of the target gene transcript and/or protein levels. Considering the concentration of siRNA in the formulation and the volume used, silencing was achieved with relatively low rates of siRNA. Further development of nanoparticle delivery methods, such as carbon dots, could be a valuable tool for basic research in plant biology and may have agricultural applications.

## EXPERIMENTAL PROCEDURES

### Plants and growth conditions

The plants used in this study included the 16C *GFP* expression line of *N. benthamiana* (Voinnet and Baulcombe, 1997) and a tomato line with constitutive expression of an *enhanced GFP* (*eGFP*). The tomato *eGFP* expression line was supplied by Seminis Vegetable. Constitutive expression of an *enhanced GFP (eGFP)* gene in tomato was accomplished with the TraitMaker ^tm^ technology developed by Mendel Biotechnology (Ratcliffe *et al*., 2008). The HP375 line of tomato was transformed with a construct expressing a LEXA DNA binding domain fused to a GAL4 activation domain under the control of a *35s* promoter. Another transgenic line was generated in the same tomato background with the *enhanced GFP* transgene and an upstream *LexA operator* (*opLEXA*) sequence. The two lines were crossed and a line that was homozygous for both insertions was selected in a subsequent generation. Transactivation of the *opLexA::eGFP* by the LEXA/GAL4 fusion protein resulted in constitutive expression of the eGFP reporter protein.

Plants were kept in a growth chamber maintained at 25°C with a light intensity of 150 μmol/m^2^/s and a day length was 16 hours. The relative humidity was not controlled and fluctuated according to irrigation frequency and plant density in the chambers at any given time. Plants were grown in a 2.5-inch pots that were irrigated by an ebb and flow system with Peters professional 20-20-20 fertilizer.

### Synthesis and purification of carbon dots from polyethyleneimines

The polyethyleneimine (PEI) used in this study included a 25 KD bPEI (Sigma-Aldrich, St. Louis, MO). Branched PEIs (bPEIs) with average Mw of 1200, 1800, and 10 KD were obtained from Polysciences (Warrington, PA). The 5 KD bPEI, Lupasol G100, from BASF was supplied as a 50% solution (w/w)—prior to carbon dot preparation the water was removed by lyophilization. Carbon dots were produced by a solvothermal reaction of the bPEIs in a solution of chloroform and methanol (4:1). Using a microwave synthesizer (Monowave 50, Anton Paar, Austria) 375 mg of bPEI in 5 mL of the solvent was heated to 155 °C with a ramp time of 5 minutes and then held at 155 °C for an additional 7 minutes. The reaction products were dried under nitrogen and then resuspended in water. Any residual chloroform was separated from the aqueous solution of carbon dots by centrifugation at 7500 g for 5 minutes. The pH of the carbon dot solution was adjusted to 8.0 with 4N HCl and the final volume was adjusted to 5 mL with water. For visible and fluorescence spectra, an aliquot was diluted 1 to 16 in water.

An alternative method was developed to produce highly efficacious carbon dots from glucose and 5 KD bPEI that does not require the use of chloroform. For the PEI functionalized glucose carbon dots, a 5 mL solution containing 37.5 mg/mL glucose and 75 mg/mL 5 KD bPEI was adjusted to pH 8.0 with 4N HCl and degassed under vacuum. The solution was then heated in a microwave synthesizer to 100 °C over 3 minutes and then held at 100 °C for an additional 5 minutes.

### FPLC fractionation of carbon dot preparations

Carbon dot preparations were purified by size exclusion chromatography. A 2.5 × 20 cm Econocolumn (Bio-Rad, Hercules, CA) filled with Sephadex G-50 superfine (GE Healthcare, Chicago, IL) was run on a Bio-Rad BioLogic Duo Flow FPLC system equipped with a QuadTec detector. The column was eluted at 2 mL/minute with 50 mM NaCl. Elution of the carbon dots was monitored at 360 nm. Five mL fractions were collected starting after 30 mL.

### RNase protection assay

For RNase protection assays, a 124 bp dsRNA was formulated with and without carbon dots in Rnase III buffer, which consisted of 20 mM Tris-HCl pH 8.0, 0.5 mM EDTA, 5 mM MgCl2, 1 mM DTT, 140 mM NaCl, 2.7 mM KCl, and 5% glycerol (w/v). Digests contained 320 ng of dsRNA and 60 ng RNase III in a total volume of 20 μL. At the indicated times, digestions were stopped with the addition of EDTA to a final concentration of 10 mM and SDS to a final concentration of 1% (w/v). Any precipitate was removed by centrifugation at 20,000 *g* for 5 minutes. The digests were run on a 1.5% TBE agarose gel.

### Formulation of carbon dots

The final concentration of siRNA in the formulations was 12 ng/μL or less. At higher concentrations siRNA aggregation can occur, which can reduce the silencing efficacy of formulations. A similar observation has been made with the carbon dot formulations used to deliver plasmid DNA to animal cells (Pierrat *et al*., 2015). The optimal concentration of carbon dots was determined empirically. Generally, a mass ratio of 40-50 for carbon dots/siRNA worked well. The siRNA and the carbon dots were added separately to two tubes containing 10 mM MES buffer pH 5.7 and 20 mM glycerol. The two tubes were then combined with vortex mixing. Formulations were then left at room temperature for at least one hour. Longer incubations at room temperature (up to a week) or at 4°C did not significantly decrease the efficacy of the formulations.

### Spray application

The spreading surfactant BREAK-THRU S279 was added to a final concentration of 0.4% within an hour of spraying. The spray application was done with an Iwata HP M1 airbrush with approximately 12 PSI and the fluid adjustment knob was set to 1.5. With a slight depression of the airbrush trigger, a very light coat of the formulation was applied to the adaxial side of *N. benthamiana* leaves or the abaxial side of tomato leaves. Efficient “flooding” of the formulation was apparent by a subtle change in the shade of green.

### Imaging and analysis

The leaves were photographed using a custom-built imaging station equipped with a Cannon EOS 70D camera with an EFS 18-55mm macro 0.25m/0.8ft lens and a high intensity blue LED light source (SL3500-D equipped with a 460 nm filter from Photon System Instruments, Czech Republic). Images were acquired using the Cannon EOS utility 2 software with tethered image acquisition. For the GFP imaging, 58 mm Tiffen Green #11 and Yellow #12 filters were used in combination to exclude wavelengths less than ~480 nm and greater than ~600 nm. An ISO of 800, a 0.25 second exposure time, and a F stop of 4.5 were typical camera settings for GFP image acquisition.

The images were processed using ImageJ (https://imagej.nih.gov/ij/index.html). Briefly, the program operator utilized the threshold color panel to highlight a border around each leaf. A border image was overlaid onto the leaf image and the pixel number within the leaf border was quantitated by the software. The quantitated number of pixels represented the total leaf area. A similar thresholding process was used to highlight a border around the GFP silencing phenotype. The GFP silenced areas were calculated by dividing the phenotypic area in pixels by the total leaf area in pixels.

### Validation of gene silencing

For qRT-PCR analysis, total RNA was extracted from phenotypic leaf tissue collected 5 days after treatment using Trizol reagent (Invitrogen) following manufacturers protocol. The RNA was dissolved in water, and the concentration was measured using Quant-iT RNA assay kit (Invitrogen, Carlsbad, CA). The total RNA samples were diluted to 5 ng/μl and 50 ng of total RNA was used to synthesize random-primed, first-strand cDNA using the High-Capacity Reverse transcription kit (Applied Biosystems, Foster city, CA). The reverse transcription products were used as template for qPCR. The qPCR reaction mixtures consisted of 2 μl cDNA, 3 μl of a primer/probe mix (0.5 μM each primer and 0.25 μM probe final concentration), and 5μl Taqman Universal PCR Master Mix (Applied Biosystems, Foster city, CA). The sequences for the primer-probe sets are provided in Table S2. The reactions were performed using an Applied Biosystems 7900HT Fast Real-Time PCR System with 40 cycles of two-step cycling at 95 °C for 15s and 60 °C for 60s. Target gene expression was expressed relative to a reference gene, *Protein Phosphatase 2a* (Nbv6.1trP16930; http://benthgenome.qut.edu.au/). Expression values were calculated using the comparative CT method: 2 ^−(Ct Target − Ct Reference)^.

For Western analysis of *N. benthamiana* or tomato, whole leaves were frozen and then ground to a fine powder. Approximately 200 mg of ground tissue was homogenized in 300 μL of ice cold buffer containing 20 mM Tris-Cl pH 8.0, 150 mM NaCl, 0.1% Triton X-100, and a protease inhibitor cocktail. The insoluble debris was removed by centrifugation. The protein concentration was quantified by Bradford assays and 10 μg of total protein for each sample was run on a 12.5% Criterion™ Tris-HCl protein gel (3450014, Bio-Rad, Hercules, CA). Following electrophoresis, proteins were transferred onto a PVDF membrane and then blocked overnight with 5% skim milk in Tris buffered saline + 0.1% Tween 20 (TBST). For GFP detection, a HRP conjugated antibody directed against the full-length GFP protein (sc-8334, Santa Cruz Biotechnology, Santa Cruz, CA) was diluted 1:1000 in TBST and 5% skim milk and incubated with the blot for 1 hour. The blot was then washed four times with TBST for 10 minutes. Pierce ECL plus western substrate (32132, Thermo Scientific Pierce Protein Biology) was used for chemiluminescent detection of GFP. Band intensity for each sample was quantified by imageJ.

## ACKNOWLEDGEMENTS

Pending permission of individuals currently on vacation

## AUTHOR CONTRIBUTIONS

SHS, BH, and WZ conceived of the experiments. BH, PH, RAS, SHS, and WZ performed the experiments. SHS wrote the original manuscript. All authors read the manuscript and assisted in the editing.

## CONFLICT OF INTEREST

The authors declare no conflict of interest.

## SUPPORTING INFORMATION

**Figure S1.**
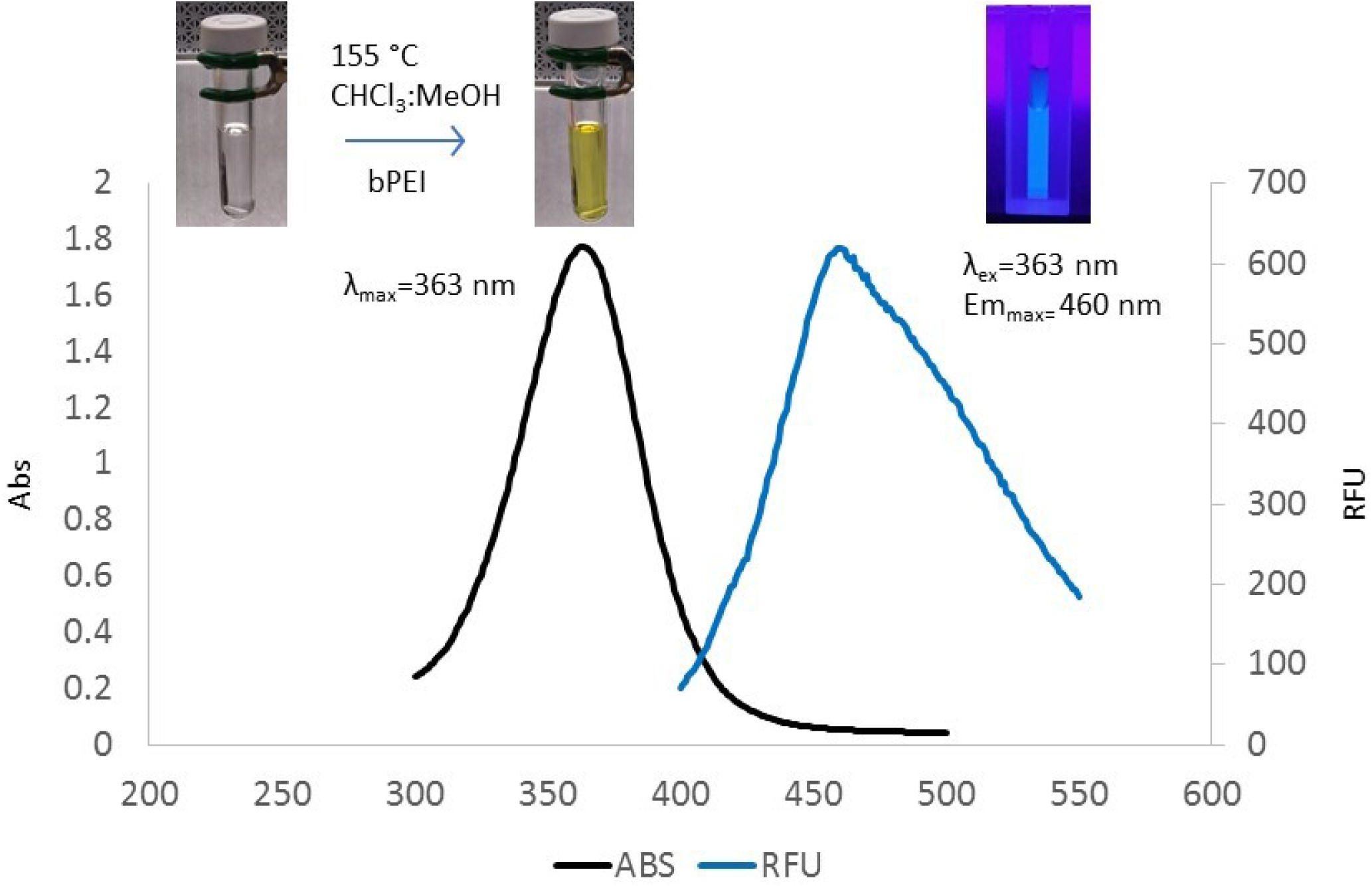
Spectroscopic characterization of carbon dots produced from polyethyleneimines.

**Figure S2.**
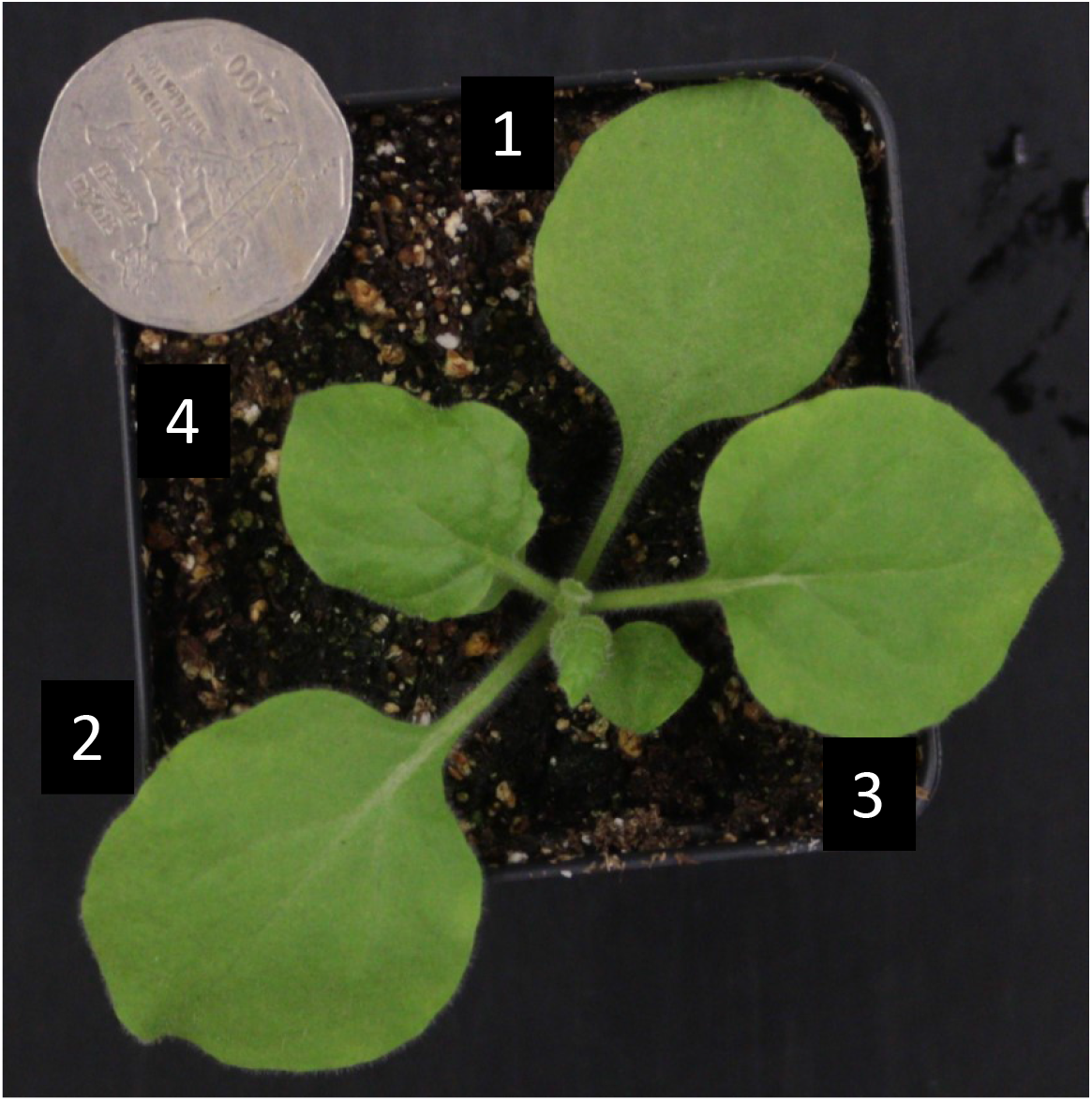
*N. benthamiana* 17 days after sowing at the time of treatment.

**Figure S3.**
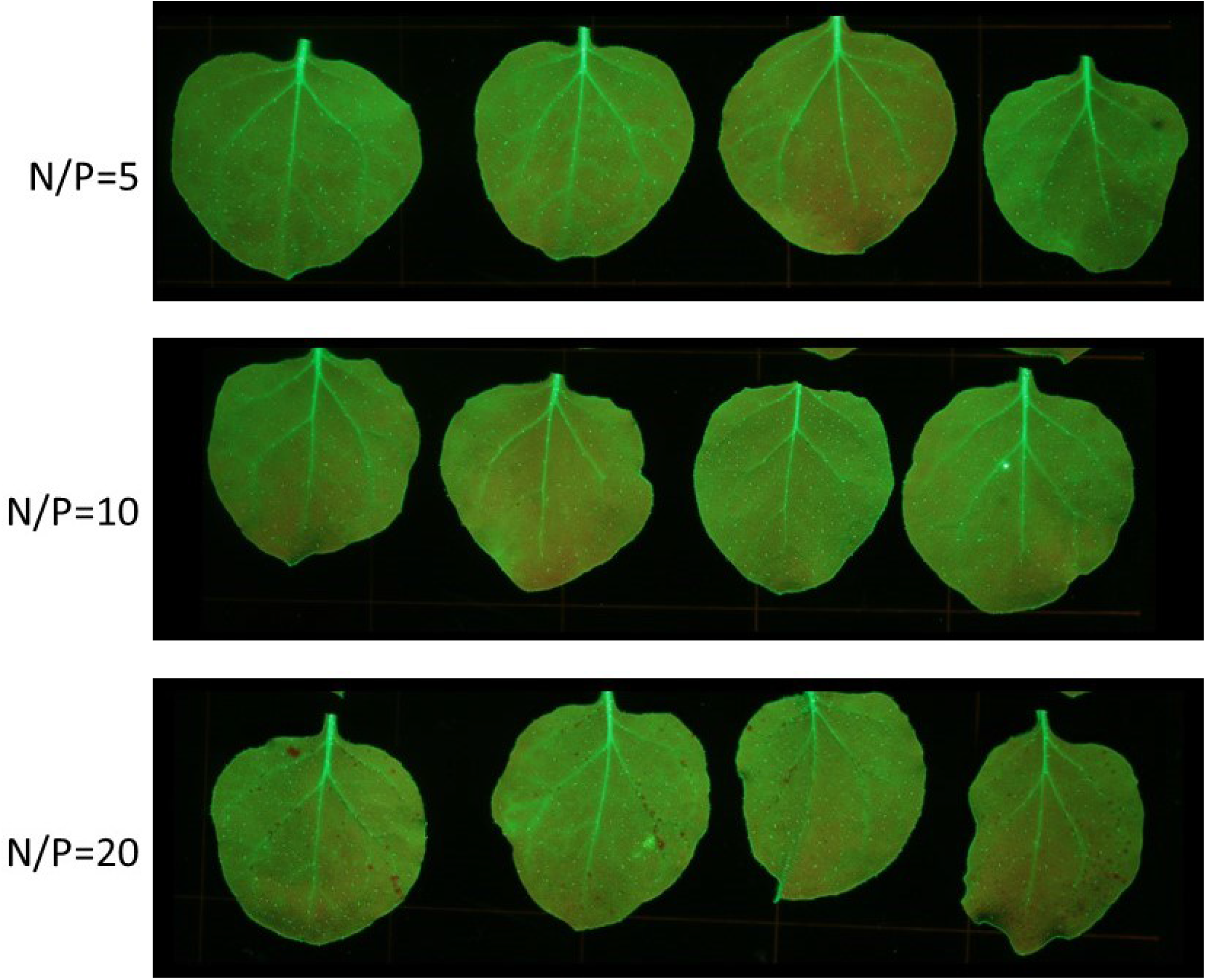
Silencing assays with unmodified 5 KD bPEI. Using the same buffer that was used for the carbon dot formulations, the PEI was tested at N/P ratios of 5, 10, and 20. Where the N/P ratio refers to the ratio of nitrogen atoms in the PEI relative to the phosphates from the nucleic acid backbone.

**Figure S4.**
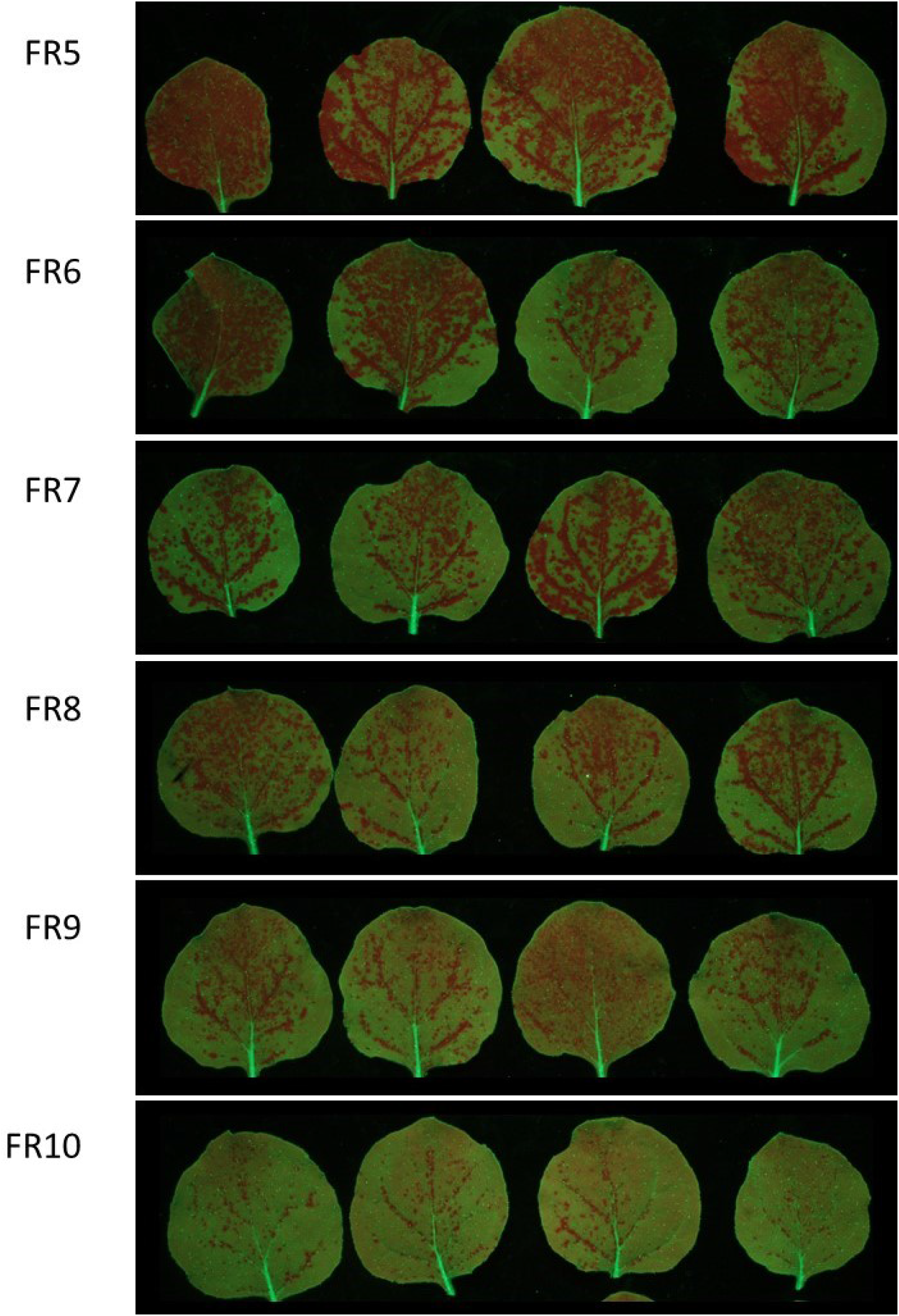
The efficacy of 5 KD bPEI (CD-5K) fractions from size exclusion chromatography. The concentration of the 22-mer targeting *GFP* was 8 ng/μL.

**Figure S5.**
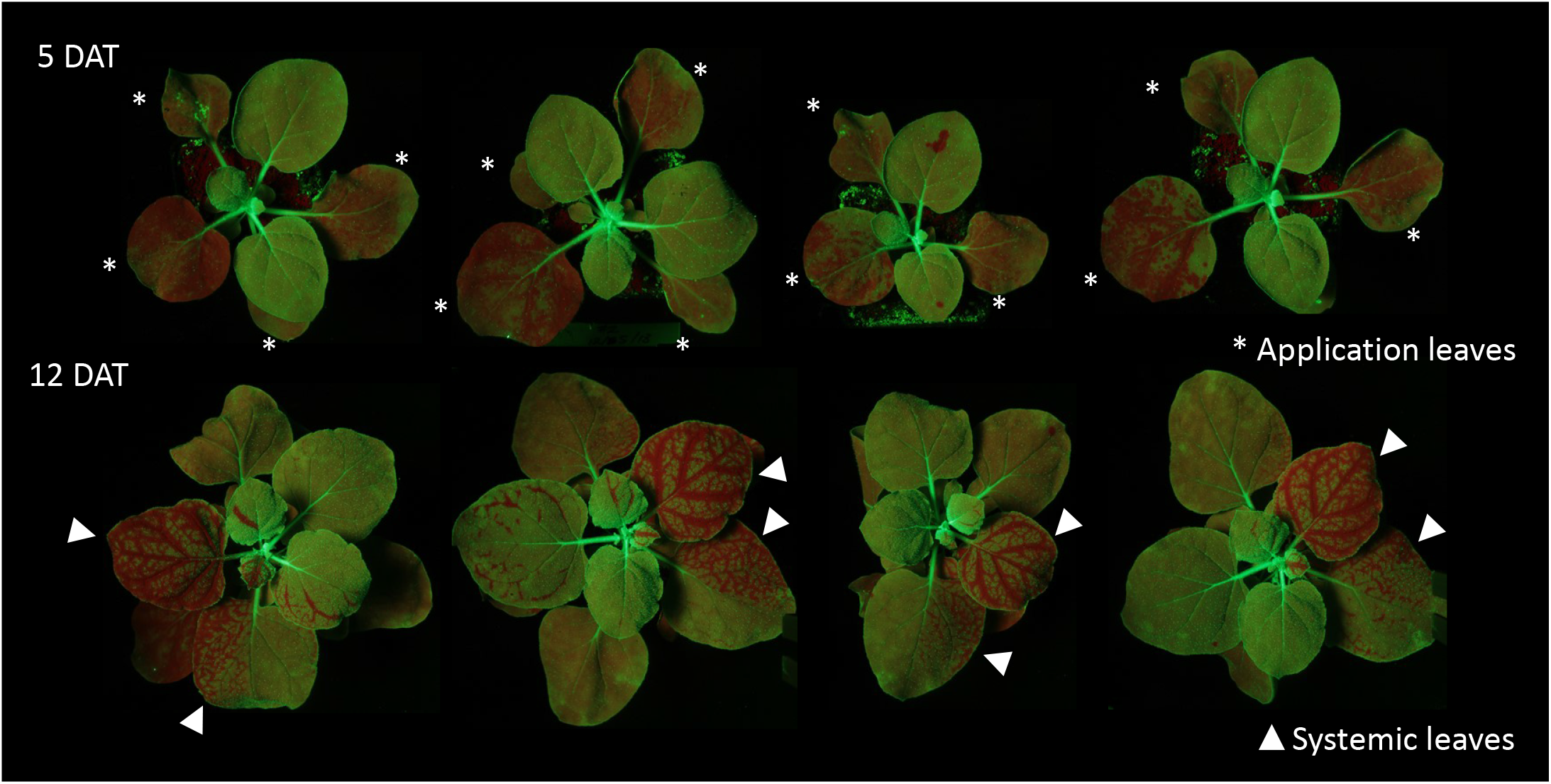
Systemic silencing of *GFP* is initiated in 16C with carbon dot formulations. *GFP* silencing on the application leaves was monitored at 5 days after treatment. Systemic spread to the newly emerging leaves was monitored at 12 days after treatment.

**Figure S6.**
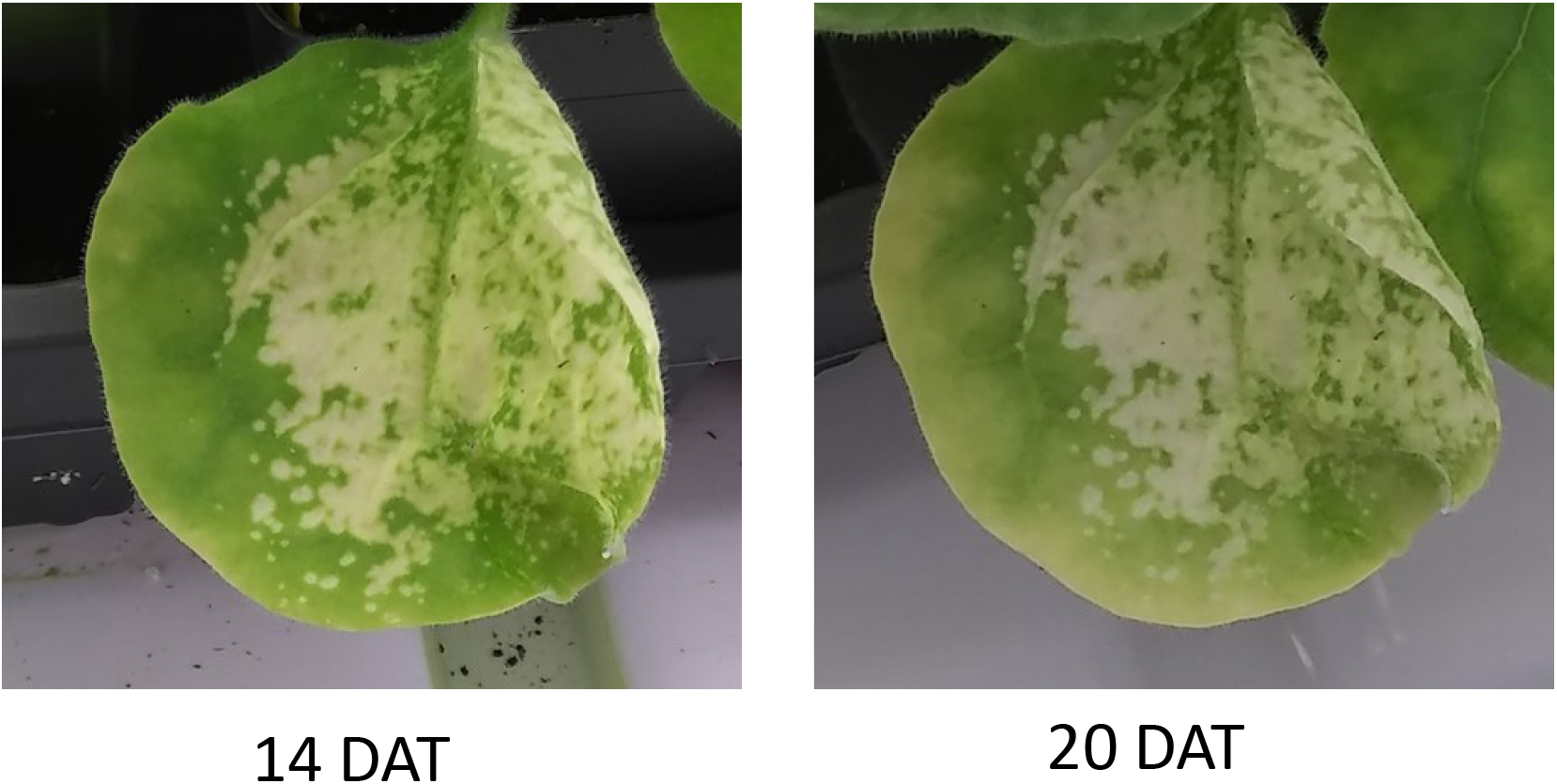
Persistence of *MgCheH* silencing phenotype in *N. benthamiana*.

**Figure S7.**
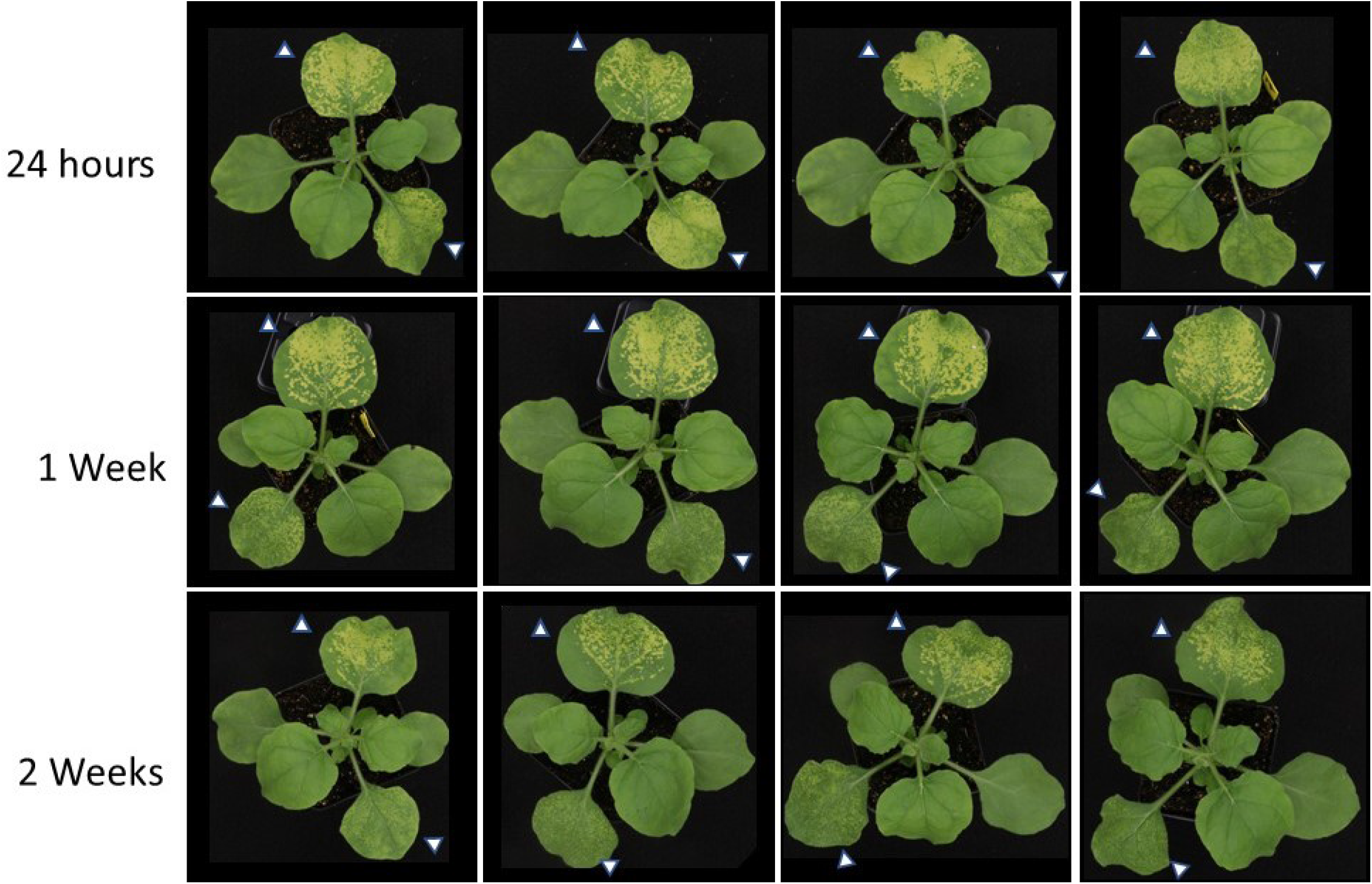
The colloidal stability and efficacy of a carbon dot formulation with the 22-mer targeting *MgCheH* was tested. The final concentration of siRNAs in the formulations was 12 ng/μL. Leaves 3 and 4 were sprayed after storage of the formulation for 24 hours, 1 week, or 2 weeks at room temperature. Plants were imaged 4 days after treatment.

**Table S1.**
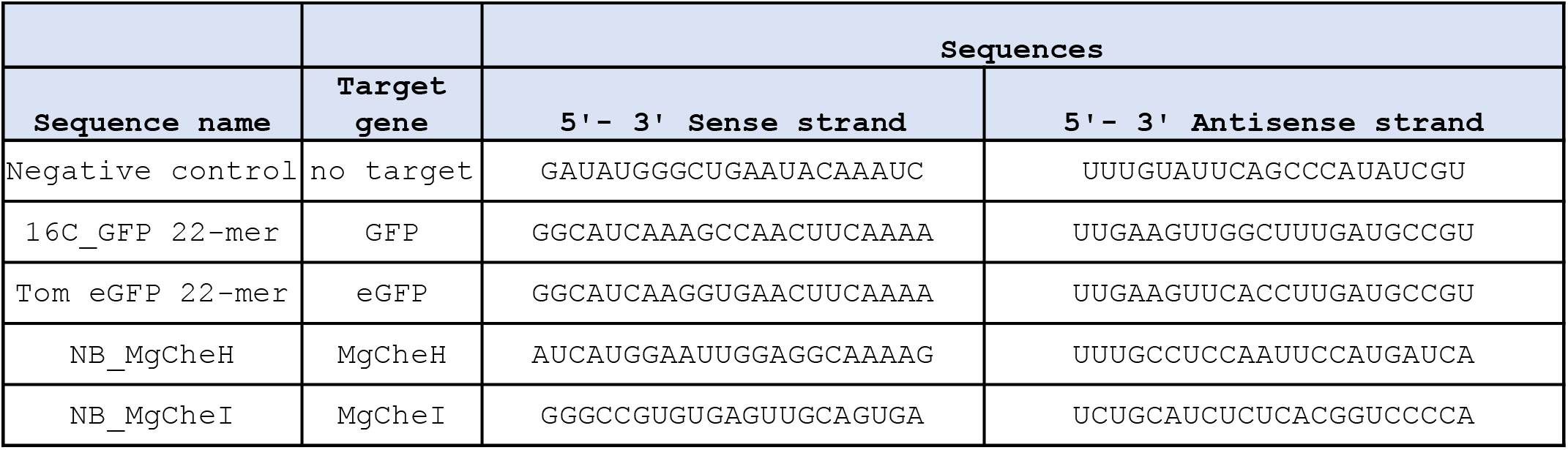
siRNA sequences used in this study. When annealed the 22-mers contained a 3’ 2-nucleotide overhang.

**Table S2.**
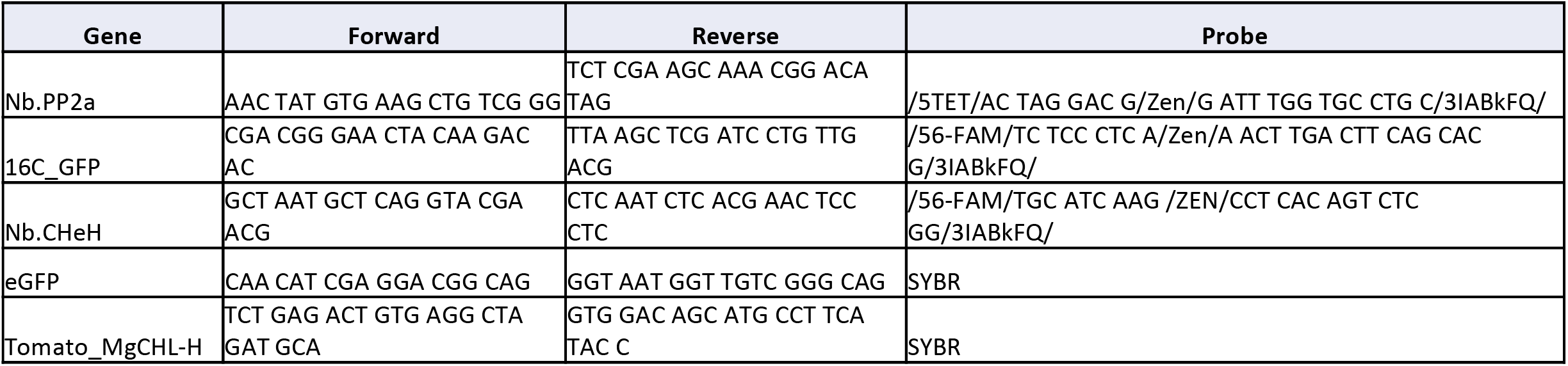
Oligonucleotide and probe sequences for qRT-PCR analysis of transcript levels.

